# ssbio: A Python Framework for Structural Systems Biology

**DOI:** 10.1101/165506

**Authors:** Nathan Mih, Elizabeth Brunk, Ke Chen, Edward Catoiu, Anand Sastry, Erol Kavvas, Jonathan M. Monk, Zhen Zhang, Bernhard O. Palsson

**Author notes:** Correspondence should be addressed to: B.O.P.

## Abstract

**Summary:** Working with protein structures at the genome-scale has been challenging in a variety of ways. Here, we present *ssbio*, a Python package that provides a framework to easily work with structural information in the context of genome-scale network reconstructions, which can contain thousands of individual proteins. The *ssbio* package provides an automated pipeline to construct high quality genome-scale models with protein structures (GEM-PROs), wrappers to popular third-party programs to compute associated protein properties, and methods to visualize and annotate structures directly in Jupyter notebooks, thus lowering the barrier of linking 3D structural data with established systems workflows.

**Availability and Implementation:** *ssbio* is implemented in Python and available to download under the MIT license at http://github.com/SBRG/ssbio. Documentation and Jupyter notebook tutorials are available at http://ssbio.readthedocs.io/en/latest/. Interactive notebooks can be launched using Binder at https://mybinder.org/v2/gh/SBRG/ssbio/master?filepath=Binder.ipynb.

**Supplementary Information:** Supplementary data are available at *Bioinformatics* online.

## Introduction

Merging the disciplines of structural and systems biology remains promising in a variety of ways, but differences in the fields present a learning curve for those looking toward this integration within their own research. Beltrao et al. stated it best, that “apparently structural biology and systems biology look like two different universes” (Beltrao et al. 2007). A great number of software tools exist within the structural bioinformatics community (Biasini et al. 2010; Grünberg et al. 2007; Gu & Bourne 2009; Hamelryck & Manderick 2003; O’Donoghue et al. 2015), and with recent advances in structure determination techniques, the number of experimental structures in the Protein Data Bank (PDB) continues to steadily rise (Mizianty et al. 2014). The challenges of integrating external data and software tools into systems analyses have been detailed (Ghosh et al. 2011), and structural information is no exception to the norm. At the systems-level, curated network models such as genome-scale metabolic models (GEMs) provide a context for molecular interactions in a functional cell (O’Brien et al. 2015). Recently, GEMs integrated with protein structures (GEM-PROs) have extended these models to explicitly utilize 3D structural data alongside modeling methods to substantiate a number of hypotheses, as we explain below.

Here, we present *ssbio*, a Python package designed with the goal of lowering the learning curve associated with efforts in structural systems biology. *ssbio* directly integrates with and builds upon the COBRApy toolkit (Ebrahim et al. 2013) allowing for seamless integration with existing GEMs. The core functionality of *ssbio* is additionally extended by hooks to many popular third-party structural bioinformatics algorithms, such as DSSP, MSMS, SCRATCH, I-TASSER, and others (see Supplementary Tables S1 and S2 for a full list) (Cheng et al. 2005; Kabsch & Sander 1983; Roy et al. 2010; Sanner et al. 1996).

## Protein class

*ssbio* adds a Protein class as an attribute to a COBRApy Gene and is representative of the gene’s translated polypeptide chain (Fig. 1A). A Protein holds related amino acid sequences and structures, and a single representative sequence and structure can be set from these. This simplifies network analyses by enabling the properties of all or a subset of proteins to be computed and subsequently queried for. For details on these properties, as well as installation and execution instructions for the third-party software used to compute them, please see the documentation. Additionally, proteins with multiple structures available in the PDB can be subjected to QC/QA based on set cutoffs such as sequence coverage and X-ray resolution. Proteins with no structures available can be prepared for homology modeling through platforms such as I-TASSER (Roy et al. 2010). Biopython representations of sequences (SeqRecord objects) and structures (Structure objects) are utilized to allow access to analysis functions available for their respective objects (Fig. 1B) (Cock et al. 2009). Finally, all information contained in a Protein (or in the context of a network model, multiple proteins) can be saved and shared as a JavaScript Object Notation (JSON) file.

**Fig. 1.**
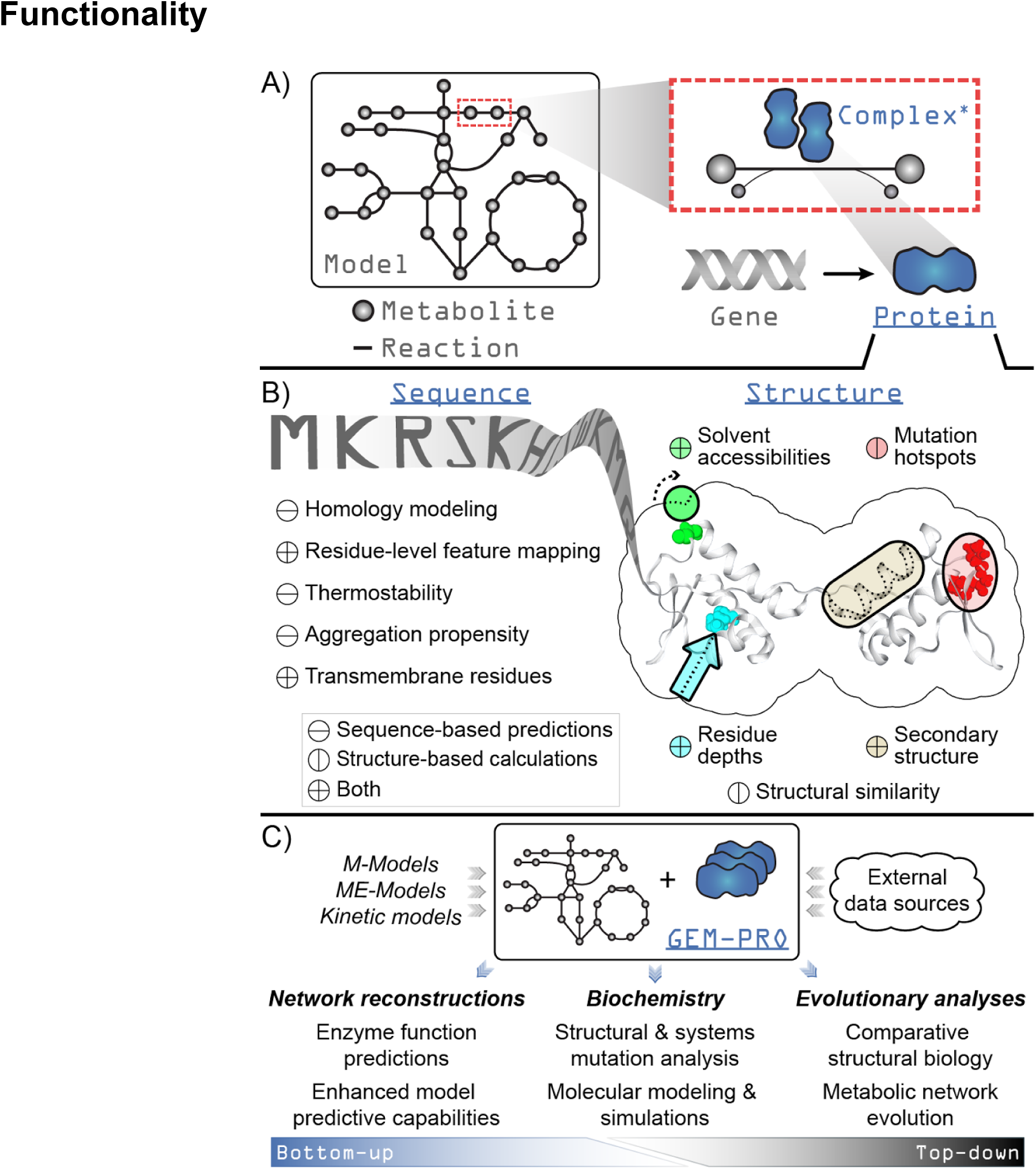
Overview of the design and functionality of *ssbio.* Underlined fixed-width text in blue indicates added functionality to COBRApy for a genome-scale model loaded using *ssbio*. A) A simplified schematic showing the addition of a Protein to the core objects of COBRApy (fixed-width text in gray). A gene is directly associated with a protein, which can act as a monomeric enzyme or form an active complex with itself or other proteins (the asterisk denotes that methods for complexes are currently under development). B) Summary of functions available for computing properties on a protein sequence or structure. C) Uses of a GEM-PRO, from the bottom-up and the top-down. Once all protein sequences and structures are mapped to a genome-scale model, the resulting GEM-PRO has uses in multiple areas of study, as noted in the main text.

## GEM-PRO pipeline

The objectives of the GEM-PRO pipeline have previously been detailed (Brunk et al. 2016). A GEM-PRO directly integrates structural information within a curated GEM (Fig. 1C), and streamlines identifier mapping, representative object selection, and property calculation for a set of proteins. The pipeline provided in *ssbio* functions with an input of a GEM, but alternatively works with a list of gene identifiers or their protein sequences if network information is unavailable.

The added context of manually curated network interactions to protein structures enables different scales of analyses. For instance, from the top-down, global non-variant properties of protein structures such as the distribution of fold types can be compared within or between organisms (Brunk et al. 2016; Monk et al. 2017; Zhang et al. 2009). From the bottom-up, structural properties predicted from sequence or calculated from structure can be utilized to guide a metabolic reconstruction (Broddrick et al. 2016) or to enhance model predictive capabilities (Chang et al. 2010, 2013; Chen et al. 2017; Mih et al. 2016). Looking forward, applications to multi-strain modelling techniques (Bosi et al. 2016; Monk et al. 2016; Ong et al. 2014) would allow strain-specific changes to be investigated at the molecular level, potentially explaining phenotypic differences or strain adaptations to certain environments.

## Scientific analysis environment

We provide a number of Jupyter notebook tutorials to demonstrate analyses at different scales (i.e. for a single protein sequence or structure, set of proteins, or network model). These notebooks can be launched in a virtual environment through the Binder project (https://mybinder.org/), with most third-party software pre-installed so users can immediately run through tutorials and experiment with them. Certain data can be represented as Pandas DataFrames (McKinney 2012), enabling quick data manipulation and graphical visualization. These notebooks are further extended by visualization tools such as NGLview for interacting with and annotating 3D structures (Nguyen et al. 2017; Rose & Hildebrand 2015), and Escher for constructing and viewing biological pathways (King et al. 2015) (Supplementary Figure S1). Module organization and directory organization for cached files is further described in the Supplementary Text.

## Conclusion

*ssbio* provides a Python framework for systems biologists to start thinking about detailed molecular interactions and how they impact their models, and enables structural biologists to scale up and apply their expertise to multiple enzymes working together in a system. Towards a vision of whole-cell *in silico* models, structural information provides invaluable molecular-level details, and integration remains crucial.

## Acknowledgements

We would like to thank Dr. Zachary King, Patrick Phaneuf, Marta Matos, and Colton Lloyd for valuable discussions in software development, and Dr. Laurence Yang, Yara Seif, JC Lachance, and Jared Broddrick for insight into desired functionalities of the package. We would also like to thank Marc Abrams for proofreading of the manuscript.

